# The carrying capacity of a fragmented landscape depends on the home-range size of a species

**DOI:** 10.1101/2025.01.09.632254

**Authors:** Henrique M. Pereira, Gretchen C. Daily

**Affiliations:** Centro de Biologia Ambiental, Faculdade de Ciências de Lisboa Edíficio C2 - Campo Grande, 1749-016 Lisboa, Portugal; Center for Conservation Biology, Stanford University Stanford, CA 94305-5020, USA

**Keywords:** Spatially explicit model, Extinction, Territoriality, Source-Sink, PVA, Fragmentation

## Abstract

Population models have not considered the problem of home-range settlement when the grain of the landscape is smaller than the home-range size. We present an individual-based model addressing this problem that combines age-structured population dynamics, optimal foraging and habitat selection. During home-range settlement each juvenile tries to maximize her fitness, which depends on the proportion of high-quality habitat in her home range. We assume that home ranges do not overlap, which can happen because the home range is defended as a territory or because individuals avoid areas used by conspecifics. We show that the population supported by the landscape at equilibrium, the carrying capacity of the landscape, decreases with the amount of low-quality habitat cover. However, this decrease is non-linear, the carrying capacity starts to decline only below a critical habitat threshold. Furthermore, when the home-range size is larger than the grain of the landscape, the carrying capacity declines faster when the habitat is fragmented. Therefore species with small home-ranges persist in instances where species with large home-ranges go deterministically extinct. Species with large population growth rates have low critical habitat sizes, and are more resilient to habitat conversion.

## INTRODUCTION

Current extinction rates are much higher than the background extinction rates in the cenozoic fossil record (Reid 1992, May et al. 1995, Pimm et al. 1995). Habitat loss is the main driver of these human-induced extinctions (Hilton-Taylor 2000, Sala et al. 2000). However, not all species are equally sensitive to changes in land use by humans (Daily et al. 2001, Owens and Bennett 2000). Therefore, a major question in Ecology today is what drives a population extinct?

Both deterministic and stochastic phenomena can cause extinction in single population models. Deterministic phenomena include changes in fecundity or survival that result in a negative population growth rate (Possingham et al. 2001). Stochastic phenomena include environmental stochasticity (Foley 1994, Hanski 1999) and demographic stochasticity (Roughgarden 1998, Caswell 2001), and are often explored in Population Viability Analysis (Beissinger and Westphal 1998). The effects of habitat loss in single population models can be modelled as a decrease in the carrying capacity of a habitat patch (Pulliam and Danielson 1991) or as a decrease of the fecundity or survival of individuals (Doak 1995). Metapopulation models go a step further and allow the study of habitat loss through the loss of habitat patches (Hanski 1999). A different mechanism is postulated in KISS models (Skellam 1951, Pereira et al. 2004), which have a source-sink structure (Pulliam 1988). In those models it is the dispersal into unsuitable habitat that drives a population to extinction, and habitat destruction increases the loss of dispersers into poor habitat.

Despite many theoretical developments in population modelling, we still have gaps on our conceptual understanding of what causes a population to decline, and ultimately to become extinct (Caughley 1994). For example, most spatially explicit models assume that individuals occupy a single point in the landscape (e.g. Flather and Bevers 2002, Pulliam et al. 1992, Wiegand et al. 1999). However, countryside landscapes, which are becoming recognized as key components of conservation efforts (Daily et al. 2001; 2003), are composed by a mixture of native and human-dominated habitats, and in those landscapes the home-ranges of individuals may include both types of habitats. Here we develop a model that combines habitat selection, optimal foraging and territory negotiation to study the dynamics of countryside populations.

Optimal foraging theory (Stephens and Krebs 1986) has provided us a framework connecting fitness with the habitat quality in the home range of an individual. On the other hand, work on habitat selection (Fretwell 1972, Rosenzweig 1991) has shown that individuals will distribute themselves in space in order to maximize their fitnesses. A more complicated problem to solve has been how individuals negotiate space. Recent theoretical developments are now providing some insight to algorithms that individuals may use to establish home ranges or territories in space (Adams 1998, Pereira et al. 2003, Stamps and Krishnan 1999). Note that in this paper we define home range as the foraging area used by an individual, and assume that home ranges of different individuals do not overlap either because they are actively defended (the traditional definition of territories), or because individuals avoid areas exploited by conspecifics.

In our model, the algorithms that individuals use to establish home-ranges and the calculation of the fitness of the home-range owner are inspired by the aforementioned body of theory. However we do not attempt to incorporate all the details associated with a traditional optimal foraging model or a territory negotiation algorithm. Instead, we attempt to develop the simplest model that allows us to study the dynamics of a population of individuals where the home-range size is greater than the grain of the landscape. This is, each home range is not a single cell in the landscape grid, but a set of cells. In order to study the effect of habitat fragmentation, we use fractal landscapes. Fractal landscapes are an excellent tool to study fragmentation because the degree of fragmentation and percentage of cover of each habitat type can be controlled independently (With 1997, Fahrig 2001, Palmer 1992).

We start by presenting the individual-based model (IBM). We then show that there is a simple equivalent analytical model for the case where the home-range size is one cell. The analytical model is similar to the source-sink model of Pulliam (1988). We show that the relationship between population size and habitat destruction is non-linear: only below a certain critical habitat threshold does the population decline. Finally we show how the carrying capacity of a landscape depends on the degree of fragmentation and the home-range size.

## THE INDIVIDUAL BASED MODEL

The IBM keeps track of two types of entities: the landscape and the female population.

The landscape is a grid of size *N* × *N* with each cell belonging to one of *n* possible habitats (see Appendix A for a description of how the fractal landscapes are generated). Each landscape cell can be in one of two states, free or occupied. Each individual in the population has two state variables, age and home range. The home range is a vector of size *T* containing the set of landscape cells that an individual occupies. *T* is assumed to be constant for a given population or species. The IBM parameters are presented in Table 1.

**Table 1:** Model parameters and typical values used in simulations of a species with large home-range size. In simulations with small home-range size two parameters change: *T* = 1 and *N* = 20.

| Symbol | Description | Value |
| --- | --- | --- |
| $N$ | Landscape size | 160 |
| $b_0$ | Maximum annual fecundity | 2 |
| $\beta$ | Age at first breeding | 1 |
| $s$ | Survival probability | 0.4 |
| $h_1$ | Affinity for habitat 1 | 1 |
| $h_2$ | Affinity for habitat 2 | 0.1 |
| $c$ | Distance weight | 0.001 |
| $T$ | Home-range size | 64 |
| $\xi$ | Initial population size | 10 |

The model starts by seeding the landscape with a population of *ξ* individuals, that have already settled their home ranges. Each simulation year consists of the following sequential steps: reproduction of individuals in breeding age, adult mortality, juvenile home range establishment, juvenile density-dependent mortality.

### Reproduction

Females reproduce only once they have age greater or equal to the breeding age *β*. The number of female juveniles produced by a breeding female follows a Poisson distribution with mean equal to her fecundity *b*. The fecundity *b* depends on the quality of the habitat available in her home range and is given by,

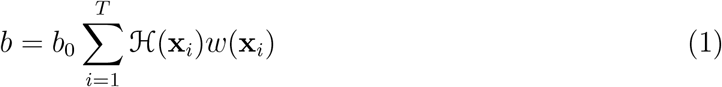

where *b*_0_ is the maximum fecundity, **x**_*i*_ = (*x, y*) is the cell *i* of the home range, ℋ(**x**) is the affinity for the habitat at position **x**, and *w*(**x**) is a weight function. The affinity function ℋ (**x**) is characteristic of each species and takes the value *h*_*i*_ for habitat type *i*, ranging from 0 (lowest quality habitat for a given species) to 1 (highest quality habitat).

The weight function gives a higher contribution to cells closer to the home range centroid 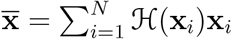, and is given by

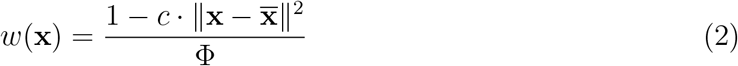

where *c* is a constant and Φ is a normalizing value so that the sum in (1) does not exceed unity,

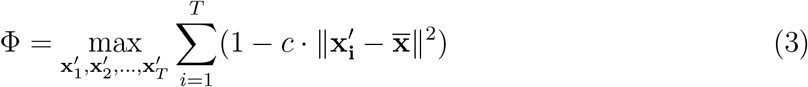

That is, Φ corresponds to a home range with a shape that maximizes fecundity. It is easy to see that this optimal shape is a home range as close to a circle as possible.

Therefore, the fecundity of an individual depends both on the quality of the habitat available within the home range and the shape of that home range, the relative importance of these two factors being determined by the constant *c*.

One can wonder what type of foraging strategies will result in fitnesses approximated by (1). For example, consider the case of a central place forager that has to carry prey to the nest. If habitat affinity is reinterpreted as the probability of prey appearing at a given cell and if the cost of travel is proportional to the square of the travelled distance, the fitness of the forager is given by (1). Note also that the optimal location for the nest is the centroid of the home range.

### Mortality

Juveniles and adults die with probability 1 − *s*. Previously occupied space is made available again.

### Juvenile home range establishment

Each juvenile tries to establish a home range so that she maximizes her fecundity, *b*. In this paper we assume global dispersal. First, the settler chooses a seed cell for her home range at random from the best habitat available in the landscape. She then iteratively adds more cells to her home range, one cell at a time, until the home range reaches size *T*. At each time in this iterative process, she analyzes all free cells that are contiguous with her growing home range, and chooses the cell that maximizes fecundity as calculated in (1). If more than one candidate cell maximize fitness, she chooses one of those cells at random. If the iterative process of home range growth stops before the home range reaches size *T* because there are no more free cells in the neighborhood of the current home range, the home range establishment process restarts with a new choice for the seed cell. The process is re-started as many times as needed until a home range of size *T* is obtained or the entire landscape has been searched. Note that the size of the home range is independent of the quality of the habitat, with the implicit assumption that the size corresponds to a foraging time constraint, i.e, it takes as much time to search for food in good habitat cells as in poor habitat cells.

This algorithm does not guarantee that an individual chooses an optimal home range. An optimal algorithm would have to look at all combinations of *T* contiguous free cells on the entire landscape, and would be very expensive computationally. Furthermore, one could argue that the search time and cognitive requirements of an optimal algorithm would be impracticable for most organisms. Instead, the algorithm used in our model gives an approximate solution to the optimal home range in the neighborhood of the seed cell.

### Juvenile density-dependent mortality

All juveniles that were not able to settle territories of size *T* are removed from the population.

### Implementation details

The individual based model was implemented as an ANSI C++ program. Our program takes as inputs the model parameters and a two dimensional matrix with the landscape. It produces a file with a matrix containing the home ranges of each individual at each time step and a matrix with the population age structure at each time step.

Our program uses the Standard C++ Library (Stroustrup 1997) extensively, which results in a high-level and compact code with only 750 lines. We use the random number generators of the library StochasticLib developed by Agner Fog. StochasticLib is freely available under the GNU licence. Our program can be downloaded from http://correio.cc.fc.ul.pt/~hpereira/software.

## THE COLONIZATION OF THE LANDSCAPE

In the remainder of this paper we consider landscapes with two habitats, with habitat 1 being preferred over habitat 2 (*h*_1_ *> h*_2_). We will refer to habitat 1 as “good habitat” and to habitat 2 as “poor habitat”. For instance, for a forest-dependent species, habitat 1 would be forest and habitat 2 could be agricultural lands. We will note the proportion of area cover by habitat *i* as *a*_*i*_. We also introduce the concept of potential carrying capacity of the landscape, *K* as the maximum number of territories that can be packed into the landscape, that is, *N* ^2^*/T*.

Consider first the case of a species with large home-range in a landscape with low fragmentation (Figure 1*A*). At time 0, all individuals can settle their home-ranges inside the large chunk of good habitat. At time 15, the good habitat is completely covered with home-ranges, and some individuals had to extend their home-ranges to poor habitat areas, due to the lack of space in the good habitat. At time 30, the population size has reached an equilibrium (Figure 2), with the good habitat completely full and some individuals settling entire home ranges in the poor habitat. This is a case of a source-sink dynamics (Pulliam 1988) with the source, the good habitat, producing enough juveniles not only to replace dead adults in the good habitat but also to colonize the sink, where very few juveniles are produced.

**Fig. 1.**
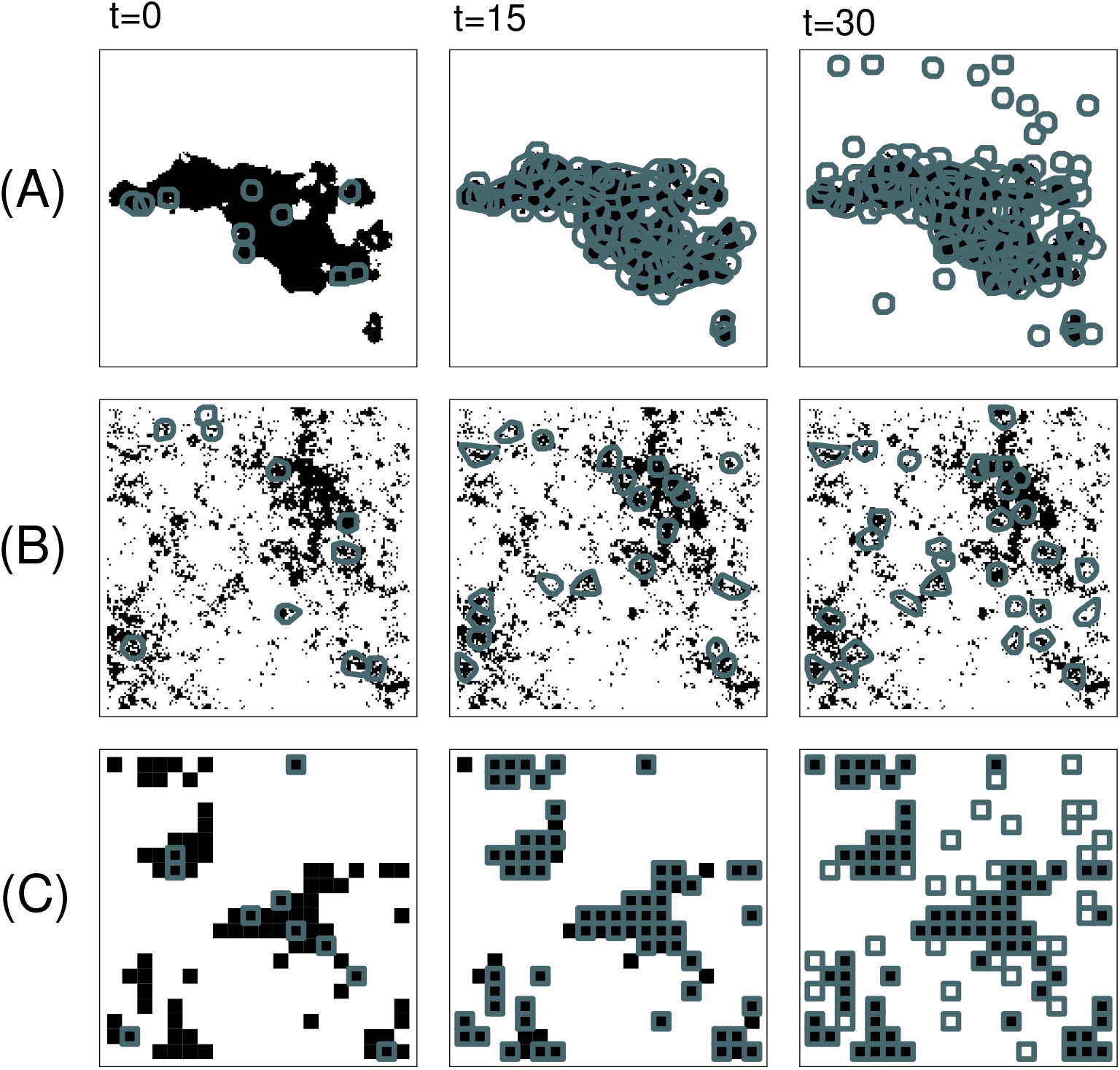
Snapshots of the home-ranges at times *t* = 0, 15 and 30 of simulations in a landscape with 20% cover of good habitat. Corresponding movies are available in the electronic appendix B. Black areas correspond to habitat cells of high-quality, white areas to habitat cells of low quality. The gray polygons are the home ranges. (*A*) Large home-range size in a landscape with low fragmentation, *D* = 2.01. (*B*) Large home-range size in a landscape with high fragmentation, *D* = 2.99. (*C*) Small home-range size in a landscape with high fragmentation. Parameters as in Table 1. The potential carrying capacity is the same for the three cases, and therefore the landscape in (c) is smaller but is shown at a higher magnification. Note that home ranges are represented with the minimum convex polygon method (White and Garrott 1990), and while the home ranges never overlap, the polygons sometimes do.

**Fig. 2.**
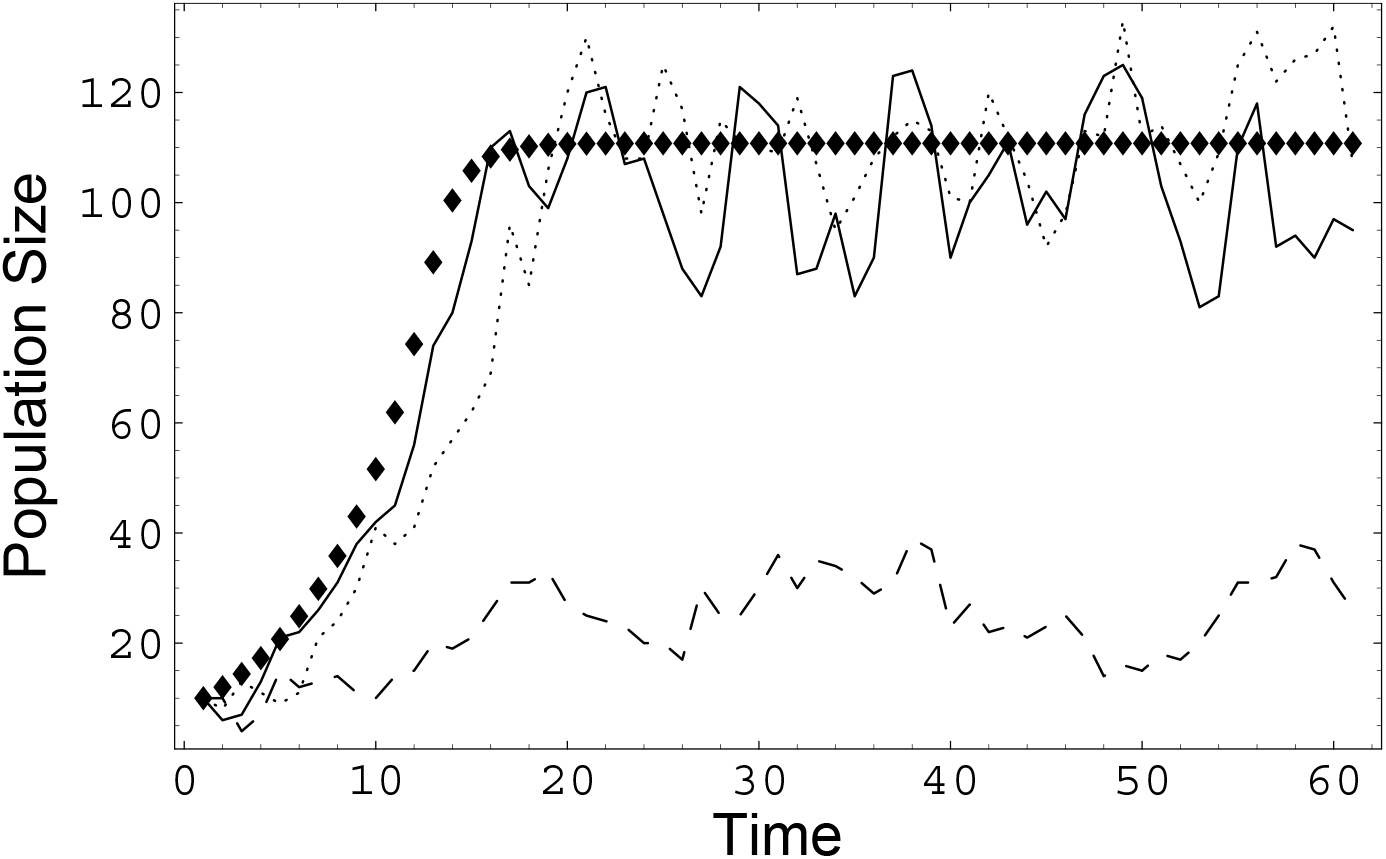
Simulations of population size as a function of time for a landscape with 20% cover of good habitat. (—) Large home-range size in a landscape with low fragmentation, *D* = 2.01. (- - -) Large home-range size in a landscape with high fragmentation, *D* = 2.99. (…) Small home-range size, fragmentation irrelevant. IBM parameters as in Table 1. Diamonds represent the prediction of the analytical model for unitary home-range size.

A very different situation arises when the landscape is highly fragmented (Figure 1*B*). Even the initial population cannot settle their home ranges entirely within the fragments of good habitat because the fragments are smaller than the home-range size (it is also important to note that the home-range settling algorithm does not perform a perfect job at finding the best home ranges). As most individuals have home ranges that include poor habitat cells, the reproductive output of the population is smaller. Therefore the population reaches an equilibrium size much smaller than in the case of the landscape with low fragmentation (Figure 2). The source-sink dynamics paradigm is not so applicable in this case, because there is not a single “large fragment” producing the juveniles in the population. Instead the landscape behaves as single unit.

Finally, consider the case of a population with unitary home-range size (Figure 1*C*). In order to study a population with the same number of individuals of the previous two cases, we reduce the landscape size (shown at a much greater magnification than the previous cases) to keep the potential carrying capacity *K* at 400 individuals. In this way, differences between simulations with large and small home-range size cannot be attributed to demographic stochasticity or a different potential carrying capacity. The landscape shown has a high degree of fragmentation, but as we shall see, that is unimportant. During the initial stages of landscape colonization (*t* = 0 and *t* = 15) all individuals are able to settle their home-ranges in the good quality habitat. This happens because the grain of the landscape is equal to the home-range size, that is, individuals can use the landscape efficiently independently of the degree of fragmentation. Only after all good habitat is full, do individuals start settling their home-ranges in poor habitat (*t* = 30). This is again a case of source-sink dynamics, but now with the source being the set of habitat patches in the landscape instead of a single patch. In effect, there is a metapopulation of good habitat patches.

It is interesting to note that the dynamics of the species with large home-range size in a landscape with low fragmentation are very similar to the dynamics of the species with small home-range size (Figure 2). That is, the populations in both cases have an exponential growth stage with the same growth rate and level off at the same equilibrium population size. We shall refer to this equilibrium as the realized carrying capacity of the landscape because it is the population that can be effectively supported by the landscape.

## ANALYTICAL MODEL FOR UNITARY HOME-RANGE SIZE

We shall now develop an analytical model for the case of unitary home-range size (*T* = 1), which will provide a theoretical background for our analysis. This analytical model is similar to the source-sink model of Pulliam (1988), but in our model the number of available home ranges in the poor habitat is limited and the breeding age may differ from one.

Let *j*(*t*) be the number of juveniles at time *t*. Let *n*_*i*_(*t, k*) be the number of individuals of age *k* with home ranges in habitat *i*. Then, the difference equations describing our model are,

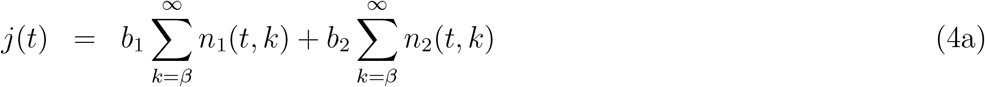

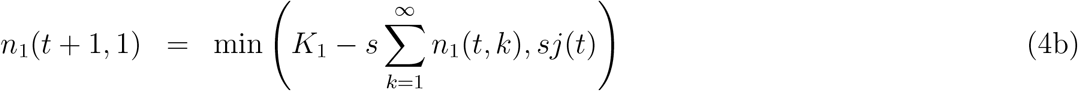

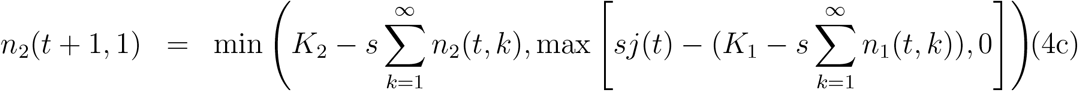

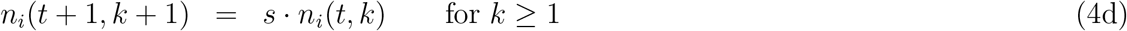

where *K*_*i*_ is the number of home range sites in habitat *i* (*K*_*i*_ = *a*_*i*_ × *N* × *N*), and *b*_*i*_ = *h*_*i*_*b*_0_ is the fecundity of females in habitat *i*. The first equation (Equation 4a) states that the number of juveniles produced each year is equal to the number of females in each habitat times the fecundities associated with each habitat. Equation (4b) states that the number of juveniles that survive the first year and settle in habitat 1 is limited by the available space. Equation (4c) states that juveniles not settling in habitat 1 will settle in habitat 2 until that habitat also reaches its potential carrying capacity *K*_2_. The last equation states that adults survive with probability *s*.

Note that for ∑ _*k*_ *n*_1_(*t, k*) ≪ *K*_1_ the population dynamics is described exactly by a Leslie matrix, and the population grows at an exponential rate *r*, which is the solution to,

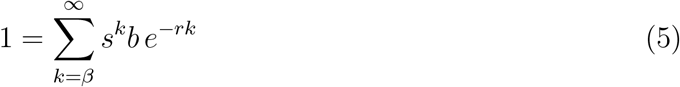

Then, as the population starts occupying habitat 2, the population growth slows down and eventually the population reaches an equilibrium population size (Figure 2).

## THE DEFORESTATION LAG

We can solve our model for the equilibrium population size when the exponential growth rate *r* is positive (otherwise the only equilibrium is zero). Let the total population size in each habitat be represented by 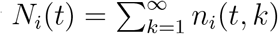. It is trivial that at equilibrium the preferred habitat is saturated, this is, 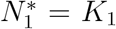. To find the population in the poor habitat we must solve,

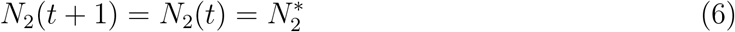

Using (4) to expand the left side of the equation we obtain,

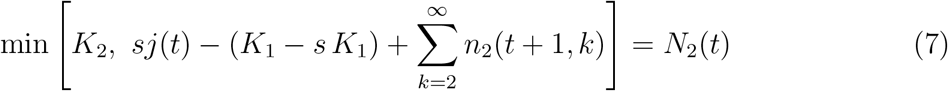

That is, either the population of the poor habitat is also at the potential carrying capacity *K*_2_, or the mortality balances recruitment at a lower carrying capacity, i.e. the realized capacity. Let us examine this second case. Substituting (4a) into (7) and noting that 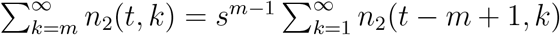 we have,

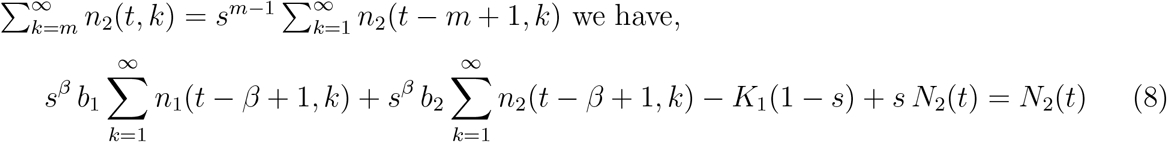

Assuming that the population was already at equilibrium at *t* − *β* + 1, we get the following simplified solution for 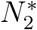,

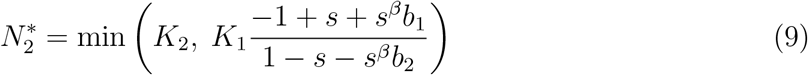

Therefore, the realized carrying capacity of the landscape 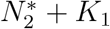, is proportional to the fecundity of females in the good habitat and inversely proportional to the fecundity of females in the poor habitat. A higher survival rate also results in a larger realized carrying capacity. A similar result had been found by Pulliam (1988).

We are interested in the case of a progressive conversion of forest habitat to non-forest habitat in a landscape of size *N* ^2^. Let *γ* be the proportion of forest converted to poor habitat. Then, we have that *K*_1_ = (1 − *γ*)*N* ^2^ and *K*_2_ = *γN* ^2^. The realized carrying capacity 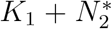 is plotted in Figure 3 along a gradient of forest loss (solid line). Initially, the loss of habitat does not have any effect on the realized carrying capacity of the landscape. This happens because the recruitment excess from the source more than compensates for the low recruitment in the sink. Beyond a critical proportion of forest loss *γ*′, the realized carrying capacity of the landscape starts declining linearly, reaching zero when no forest remains. We call this non-linear effect the deforestation lag: there is a time lag between the beginning of deforestation and the first effect on the population size of a forest-dependent species. Note that the realized capacity calculated from the analytical model is in close agreement with simulations from the IBM (dashed line).

**Fig. 3.**
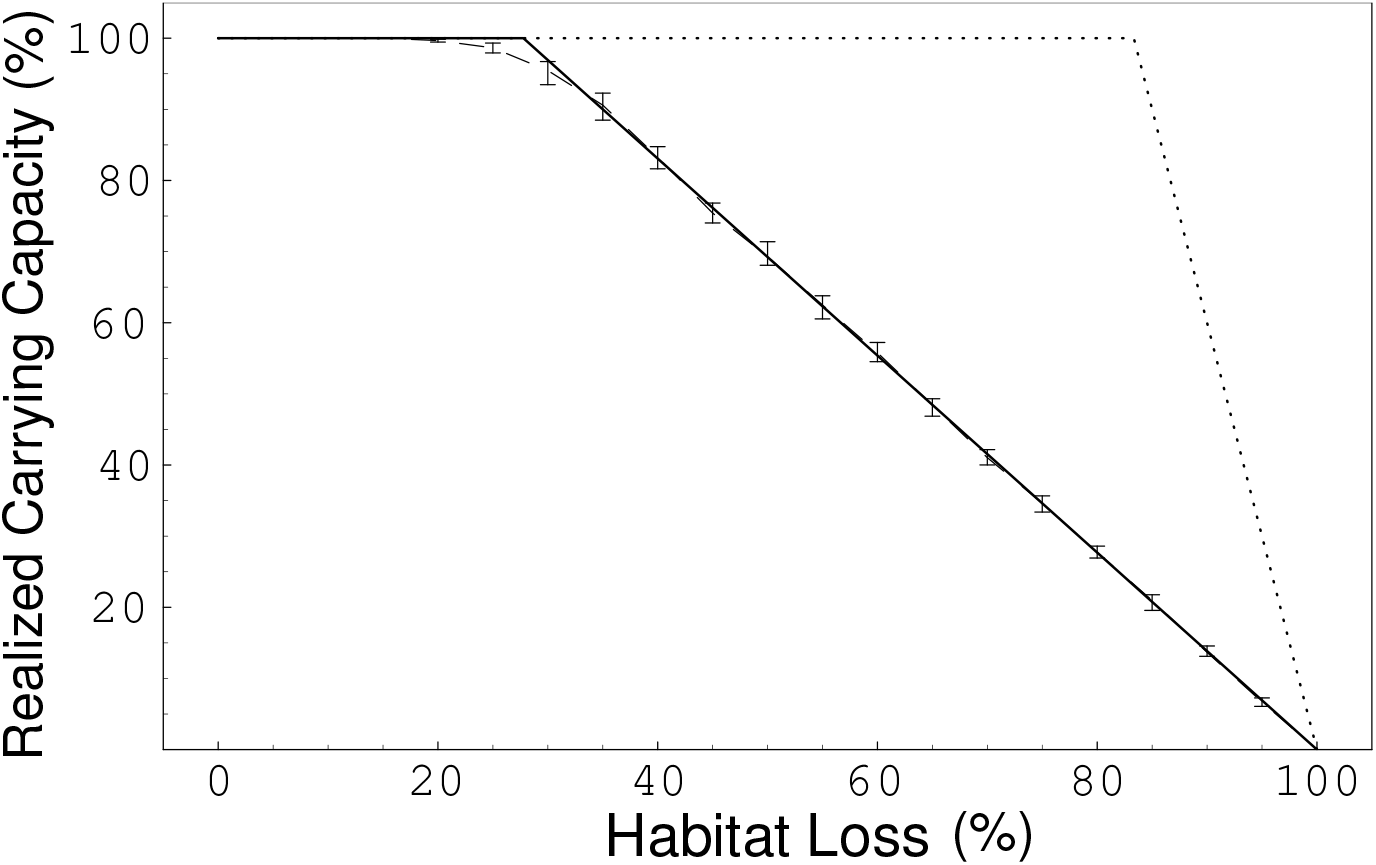
Realized carrying capacity as a function of forest loss for a species with small home-range size. (—) Analytical result. (- - -) Results from IBM simulations. By time 50 the population had reached an equilibrium with stochastic fluctuations (see Figure 2), therefore for each simulation the mean population size from time 50 to time 60 was calculated. The median of 40 such simulations is given at each point. Error bars give the interquartile intervals. (…) Species with a maximum fecundity three times greater, *b*_0_ = 6. Parameters as in Table 1.

The deforestation lag *γ*′ is in general greatest in species with large population growth rate (Figure 4), as it would be expected from (9). Species with small growth rate will start declining almost as soon as the deforestation process starts, while species with very large population growth rate starts declining only when almost no forest remains, or may not decline at all. This last case happens when the growth rate is so high, that even when the fecundity is reduced tenfold, as it happens in the poor habitat for this species (Table 1), the realized population growth rate is positive.

**Fig. 4.**
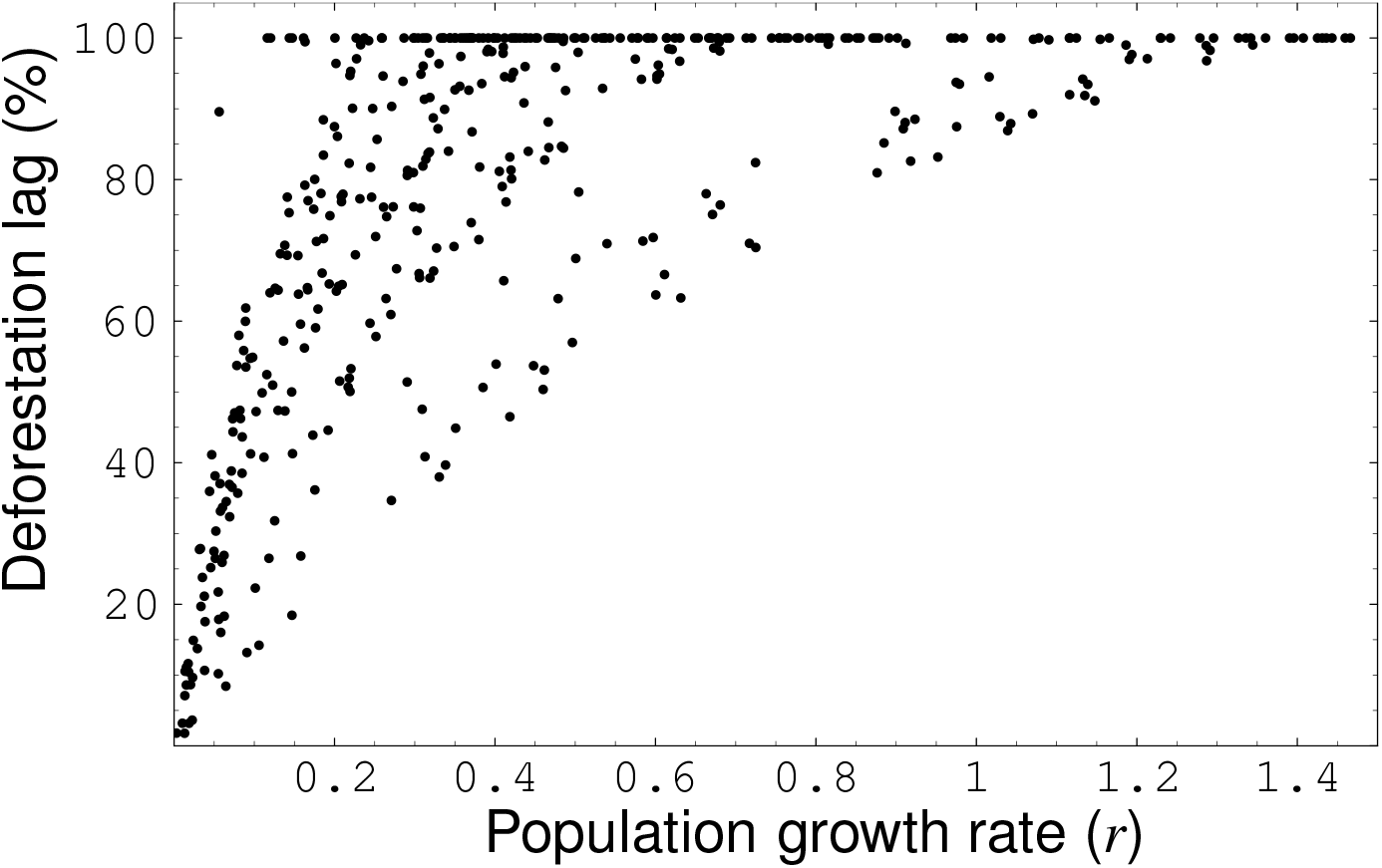
Deforestation lag as a function of population growth rate for 1000 random species. Each species corresponds to a set of life-history parameters drawn from a random distribution, as follows: *b*_0_ ~ Uniform(0, 5), *s* ~ Uniform(0, 1), *β* ~ DiscreteUniform(1, 5). Note that some species have such high growth rates that they can survive even when there is no forest left (*γ*′ = 1).

It is important to note that the decline is much faster in species where the deforestation lag is larger (Figure 3, dotted line), a phenomenon that could surprise a manager.

## THE CARRYING CAPACITY OF A FRAGMENTED LANDSCAPE

We now examine the relationship between deforestation and realized carrying capacity for different degrees of habitat fragmentation. Note that in our model, habitat fragmentation affects only species with home-range size larger than one cell, so in this section we study a species with large home-range size.

Consider first a landscape with low fragmentation (Figure 5, solid line). The pattern is very similar to the one found for unitary home-range size. In the first stages of habitat loss no significant effect occurs. Then, after a critical amount of forest loss, the realized carrying capacity starts declining in a linear fashion until reaching zero when no forest remains.

**Fig. 5.**
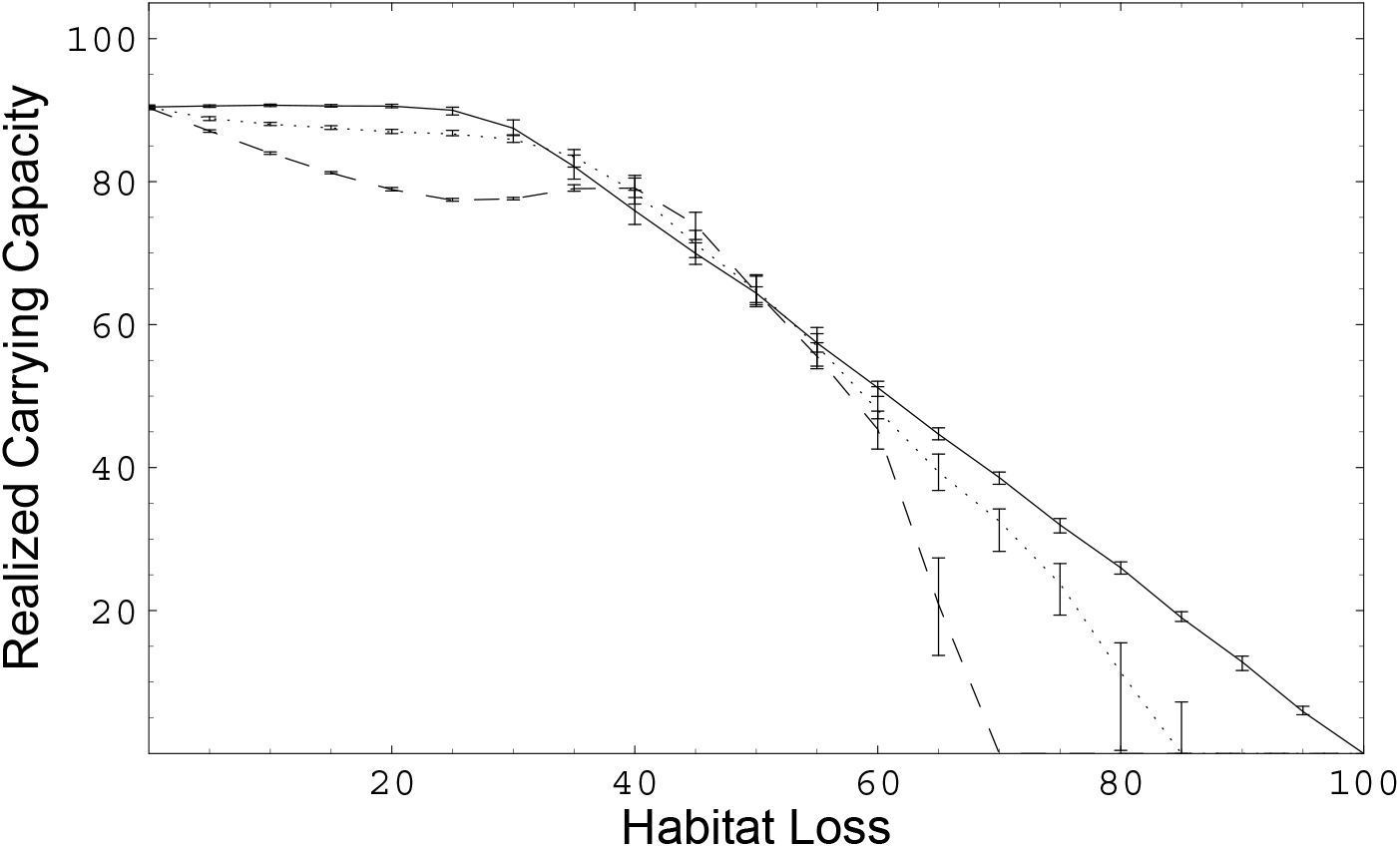
The effect of habitat fragmentation on the realized carrying capacity of a species with large home-range size. (—) Landscape with low fragmentation, *D* = 2.01. (…) Landscape with high fragmentation, *D* = 2.99. (- - -) Random landscape with no spatial auto-correlation. In each IBM simulation the mean population size from time 50 to time 60 was calculated. The median of 40 such simulations is given at each point. Error bars give the interquartile intervals. IBM parameters as in Table 1.

When the degree of fragmentation is high, a different pattern arises (Figure 5, dotted line). The decline rate of the realized carrying capacity accelerates in the later stages of forest loss, and the population goes extinct when a portion of the original forest still remains. In the case of the parameters used in this example (Table 1), the extinction occurs when 15% of the forest remains. This means that it is possible that a species with large home-range size goes extinct in a situation where a species with small home-range size persists (compare with Figure 3), despite other life-history parameter being similar between those two species. Therefore the realized carrying capacity of a landscape depends on the home-range size of a species.

The fractal landscape with D=2.99 is not the most fragmented landscape possible. A random landscape, where each point is independent of the others, has even a higher degree of fragmentation (With 2002). For such a landscape, a species with large home-range crashes even earlier in the deforestation process, when about 30% of the original forest still remains (Figure 5, dashed line). Therefore the largest the degree of fragmentation, the likeliest is the extinction of a species with large home range.

Interestingly, even when all habitat is forest (Figure 5), the realized carrying capacity is not 100%. This happens because there is always some unused space between territories that is too small for an individual to settle her home range there. Therefore, the total number of individuals in the landscape is slightly lower than if a perfect packing of territories was reached. A perfect packing of territories could be achieved for instance if territories were hexagonal, but a fixed shape would not allow an individual to shape her home range to the spatial distribution of the habitat.

## DISCUSSION

Several models have shown that the relationship between habitat loss and population abundance is non-linear. For instance, in the Levins metapopulation model a species may go extinct before all habitat patches are lost (Nee and May 1992, Tilman 1997). A similar phenomenon is apparent on the incidence function metapopulation model (Hanski 1999). What is particularly interesting with our model is that during the first stages of habitat destruction, there is no visible effect on the population density. This is in part a consequence of having a territorial model: more individuals are born than are territories available. Lande’s (1987) generalization of the Levins model to a territorial population also displays something akin to a deforestation lag, although in a smaller region of the parameter space. On the other hand, a major difference between our approach and classical metapopulation models is that we consider the whole landscape. In metapopulation models the focus is on the populations of the patches. However, many species do occupy different habitats in the landscape with different degrees of preference (Rosenzweig 1991, Milinski and Parker 1991). Therefore, habitat conversion does not mean that the new habitat becomes immediately empty. It means that individuals occupying the new habitats may obtain smaller fitnesses (or the reverse in the case of higher affinities for the new habitat). When the area of the poor habitats is large enough that recruits are insufficient to occupy the entire landscape, the effect of habitat loss becomes apparent.

The flux of population from source to sink habitats exhibits another interesting property: the majority of the population can occur in the sink habitat. This phenomenon was studied by Pulliam and colleagues (1988, 1991), who showed that e.g. 90% of the population may occur in the poor habitats, potentially misleading a manager about the importance of the good habitats holding the other 10% of the population. In our model, this situation occurs when the deforestation lag is very large.

Foley (1994) showed that the time to extinction in a stochastic environment declines with increasing carrying capacity of the population. Furthermore, the risk of extinction due to demographic stochasticity declines exponentially with increasing population size (Caswell 2001). The carrying capacity of a landscape is in general inversely proportional to home-range size. Therefore, species with large home-ranges are more vulnerable to habitat loss due to demographic and environmental stochasticity. Here, we show that species with large home-ranges are also more vulnerable to habitat loss deterministically. In our model the environment is deterministic and we control for the effect of demographic stochasticity by using the same potential carrying capacity in the simulations with large and small-home ranges. Nevertheless, our models shows that in a fragmented landscape, species with small home-range go extinct only when there is no good habitat left, while species with large home-range go extinct much earlier in the deforestation process. This happens because species with small home-range can better adapt the shape of their home-ranges to a fragmented habitat then species with large home-range size.

Some studies have found that species with large body size are particularly vulnerable to habitat loss (Gaston and Blackburn 1996, Gillespie 2001 but see Davies et al. 2000, Owens and Bennett 2000). Body mass is inversely proportional to population density (Calder 1984). Therefore, it is possible to infer that species with large-home ranges sizes are more vulnerable to habitat loss as theoretical models predict. An interesting question is which of the three factors mentioned above, demographic stochasticity, environmental stochasticity, or foraging efficiency, plays the most important role in this vulnerability.

In the standard logistic model the population growth rate *r* and the carrying capacity *K* are independent parameters. Little theoretical attention has been devoted to developing mechanistic models that relate these parameters (Kuno 1991), despite empirical evidence for a positive correlation between these parameters (Holt et al. 1997). In our model, the realized carrying capacity of the landscape is determined mechanistically by the life-history parameters of a species and by the proportion of cover of the different habitat types. In general, the greater the population growth rate, the larger the deforestation lag, resulting in a larger realized carrying capacity, unless the poor habitat is already saturated.

In our model dispersal is global and individuals are choosy. One would expect that the realized carrying capacity of the landscape would be smaller if dispersal was local, because individuals would be restricted to the set of connected patches. The picture gets more complicated when one adds error to the habitat choice process. For instance, in the Skellam (1951) model, individuals disperse at random according to a gaussian distribution. In this case, a species with larger mean dispersal distance is more vulnerable to extinction. The IBM framework presented in this model could be used to study the effect of dispersal distance and habitat choice in a systematic way.

Finally, we hope that our model provides some insights into landscape population dynamics that might be useful to managers. First, one may not perceive that a species is affected by habitat loss until late in the deforestation process. Second, what can look initially as a linear decline of the population size with the amount of habitat converted, can become a sudden crash of population size.

## APENDIX A. Generating Fractal Landscapes

First we generate a 3D random fractal of size 2*N* × 2*N* using the fourier filtering method (Saupe 1988). Briefly, this method consists of generating a spectrum corresponding to the following spectral density function,

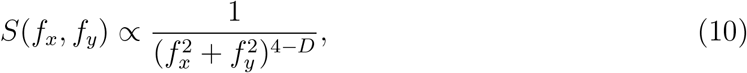

where *f*_*x*_ and *f*_*y*_ are the frequencies along the x-axis and y-axis, and *D* is the fractal dimension ranging from 2 (planar surface) to 3 (fractal filling the entire three-dimensional space). The phases at each frequency are drawn from a uniform random distribution between 0 and 2*π*. The 3D random fractal is obtained by applying the inverse fourier transform to that spectrum. In order to avoid the periodicity of the fourier transform, we use only the upper left quadrant of the random fractal, this is points *j, k* = 0 … *N* − 1, so that the final landscape has size *N* × *N*.

We wish to generate a fractal landscape with proportions of habitat cover *a*_1_, *a*_2_, …, *a*_*n*_. This is done by attributing to habitat *i* all the points of the random fractal with height between *a*_*i*−1_ and *a*_*i*_, with *z*_0_ being the lowest point of the random fractal and *a*_*i*_ being the point corresponding to the height percentile 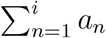. Note that this is equivalent to creating a topographic map of the landscape, and determining that the borders of habitat *i* are the altitude contour lines *a*_*i*−1_ and *a*_*i*_. For instance, consider the case of two habitats and 10% cover of habitat 1. Then *a*_1_ is the height below which 10% of the area of the random fractal is comprised. All the points below *a*_1_ belong to habitat 1 and all the points above *a*_1_ to habitat 2. Note that for the case of *n* habitats we always have that habitat *i* is contiguous to habitats *i* + 1 and *i* − 1.

A Mathematica (Wolfram 1996) package that implements this algorithm is available at: http://correio.cc.fc.ul.pt/~hpereira/software.

## ONLINE APENDIX B. Movies of the simulations in Fig. 1

Movie 1*A*: Large home-range size in a landscape with low fragmentation.

Movie 1*B* : Large home-range size in a landscape with high fragmentation.

Movie 1*C* : Small home-range size in a landscape with high fragmentation.

